# Sleep spindle frequency: overnight dynamics, afternoon nap effects, and possible circadian modulation

**DOI:** 10.1101/2021.06.28.450146

**Authors:** Róbert Bódizs, Csenge G. Horváth, Orsolya Szalárdy, Péter P. Ujma, Péter Simor, Ferenc Gombos, Ilona Kovács, Lisa Genzel, Martin Dresler

## Abstract

Homeostatic and circadian processes play a pivotal role in determining sleep structure, timing and quality. In sharp contrast with the wide accessibility of the EEG index of sleep homeostasis, an electrophysological measure of the circadian modulation of sleep is still non-available. Evidence suggests that sleep spindle frequencies decelerate during biological night. In order to test the feasibility of measuring this marker in common polysomnographic protocols, the Budapest-Munich database of sleep records (N = 251 healthy subjects, 122 females, age range: 4–69 years), as well as an afternoon nap sleep record database (N = 112 healthy subjects, 30 females, age range: 18–30 years) were analysed by the Individual Adjustment Method of sleep spindle analysis. Slow and fast sleep spindle frequencies were characterized by U-shaped overnight dynamics, with highest values in the first and the fourth-to-fifth sleep cycle and the lowest values in the middle of the sleeping period (cycles 2–3). Age-related attenuation of sleep spindle deceleration was evident. Estimated phases of the nadirs in sleep spindle frequencies were advanced in children as compared to other age groups. Additionally, nap sleep spindles were faster than night sleep spindles (0.57 and 0.39 Hz difference for slow and fast types, respectively). The fine frequency resolution analysis of sleep spindles is a feasible method of measuring the assumed circadian modulation of sleep. Moreover, age-related attenuation of circadian sleep modulation might be measurable by assessing the overnight dynamics in sleep spindle frequency. Phase of the minimal sleep spindle frequency is a putative biomarker of chronotype.

## 1 INTRODUCTION

Easily accessible and reliable biomarkers of sleep regulatory processes are of utmost importance in the objective measurement of sleep quality and rest-activity rhythms. The two-process model is one of the most influential and substantive theories of sleep regulation, proposing a linear interaction between sleep homeostasis (process S) and the circadian rhythm (process C) in humans (Borbély, 1982). Although the basic tenets and the main predictions of the model are widely accepted and empirically supported (Borbély, Daan, Wirz-Justice, & Deboer, 2016), investigators conducting common polysomnography studies are often challenged by the task of depicting and characterizing the two sleep regulatory processes. Thanks to widely available digital electroencephalogram (EEG) recording and analysis tools, sleep homeostasis is easily measurable by quantifying EEG slow wave activity (SWA: spectral power in the 0.75–4.5 Hz range) (Achermann, 2009). In contrast, the circadian modulation of sleep EEG is either assumed and hypothetically included in the model, without considering the individual differences in phase (Achermann & Borbély, 2003; Borbély, 1982) or measured in particularly complex chronobiology study settings, like constant routine (Knoblauch et al., 2005) or forced desynchrony protocols (Wei, Riel, Czeisler, & Dijk, 1999), which are not easy to implement in common clinical and research sleep studies and require notoriously large time and labor investments. Another objective method which is instrumental in the assessment of the circadian modulation of sleep consists of long-term physiological measurements of core body temperature and/or specific endocrine factors, like melatonin or cortisol release (Oster et al., 2017; Reid, 2019). Although these more direct ways of assessing the circadian component of sleep regulation seem promising, they are widely non-convenient and expensive, thus rarely included in the routine polysomnography examination protocols. Omitting these measurements leads to a permanent lack of information on the individual differences in chronotype, the latter being defined as the phase of entrainment of the circadian rhythm and the zeitgebers/local time (Roenneberg, 2012). Standard polysomnography examinations disregard the fact that subjects with earlier or later chronotypes are characterized by different levels of circadian modulation of their sleep during the same period of recording. Known age effects of chronotype include a progressive phase delay from childhood till the end of adolescence, followed by a slow gradual phase advancement during aging (Roenneberg et al., 2004). As a consequence, a widely accessible EEG index of the circadian processes could significantly improve the insight into sleep regulation and complete our understanding of sleep and chronotype.

Given the intermingling of sleep homeostasis and circadian modulation in natural, night-time sleep, specific EEG-measures of the circadian process would be advantageous. Results of studies implementing forced desynchrony, constant routine, sleep displacement or overnight polysomnography protocols suggest that various aspects of sleep spindles reflect the circadian modulation of sleep or time-of-day (Aeschbach, Dijk, & Borbély, 1997; Knoblauch et al., 2005; Purcell et al., 2017; Wei et al., 1999). Sleep spindles are known as trains of distinct sinusoidal EEG waves with a frequency of 11–16 Hz (most frequently 12–14 Hz) lasting at least 0.5 s and emerging in non-rapid-eye-movement (NREM) sleep stages N2 and N3 (Berry et al. 2018). These oscillatory events were shown to arise from the hyperpolarization-rebound sequences of widely synchronized thalamocortical neurons. The thalamic reticular nucleus is hypothesized to be the main source of hyperpolarization, whereas the T-type Ca^2+^-channels are sources of the rhythmic recurrence of firing (Fernandez & Lüthi, 2020). Based on topography and dominant frequency, two types of sleep spindles are distinguishable. The anterior (frontal) slow spindles are known to consist of waves of roughly 12 Hz (<12.5 Hz), whereas the posterior (centroparietal) fast sleep spindles are oscillations with a typical frequency of 14 Hz (>12.5 Hz) (Gibbs & Gibbs, 1951). Thorough analyses of sleep spindles indicate considerable individual differences (Bódizs, Körmendi, Rigó, & Lázár, 2009; Cox, Schapiro, Manoach, & Stickgold, 2017), as well as age and sex effects. The latter two consists of unusually low frequency sleep spindles in prepubertal (and pre-schooler) ages (Ujma, Sándor, Szakadát, Gombos, & Bódizs, 2016), as well as a slightly (0.5 Hz) increased sleep spindle frequency and variability in pubertal girls and adult women, as compared to boys and men (Bódizs et al., 2021; Ujma et al., 2014). Spindle wave frequency increases with remarkable linearity across the age range of 6–18 years (Zhang, Campbell, Dhayagude, Espino, & Feinberg, 2021), reaching a plateau in adulthood (Purcell et al., 2017), which was hypothesized to reflect the maturation of thalamocortical circuits via myelination (Zhang et al., 2021). Increased variability of sleep spindles in women is known to reflect the neural effects of hormonal variation during the menstrual cycles (Ishizuka et al., 1994). That is, the differentiation of slow and fast sleep spindles and the individual adjustment of sleep spindle frequencies is a basic requirement in conducting studies in the field. Furthermore, sleep spindles were shown to contribute to sleep maintenance, and memory consolidation, as well as to correlate with psychometric measures of intelligence (Fernandez & Lüthi, 2020; Ujma, Bódizs, & Dresler, 2020), but none of these findings were shown to be tightly associated with the small differences in oscillatory wave frequencies.

Circadian modulation of sleep spindle frequency was evidenced in a forced desynchrony study, which is instrumental in differentiating sleep homeostatic and circadian effects by invoking the non-24-hours (usually 28 hours) days approach. The frequency of sleep spindles was lowest at the nadir of the core body temperature rhythm and peaked at the acrophase of the body temperature rhythm. In addition, circadian modulation of sleep spindle frequency, that is the decrease in oscillatory frequency during the biological night, was attenuated in aged subjects, as compared to young ones (Wei et al., 1999). These findings cohere with outcomes of nap studies performed in constant routine conditions. The nadir in sleep spindle frequency was shown to coincide with the acrophase of salivary melatonin levels in humans in this report (Knoblauch et al., 2005). In addition, experimental manipulation of sleep timing indicates prominent time of day-effects in sleep spindle frequency activities, suggesting a frequency-specific circadian modulation: lower bins (12.25–13 Hz) of the spindle frequency spectral power peak during the middle of the night sleep period (between 2:00 and 5:00 AM), when the highest bins (14.25–15 Hz) reach their nadir (Aeschbach et al., 1997). In addition, daytime recovery sleep after 25 hours of wakefulness was shown to be characterized by increased spindle frequency compared to baseline night sleep records (Rosinvil et al., 2015). This latter effect was more pronounced in young as compared to middle-aged participants. That is sleep spindles are slower in the middle of the habitual sleep periods, characterized by high melatonin and low core body temperature levels. It has to be noted that brain temperature per se is a direct modulator of sleep spindle frequency: higher temperatures imply faster spindles (Csernai et al., 2019).

The above studies were not designed to differentiate between slow and fast sleep spindles, thus authors could not deduce whether the reported post-midnight deceleration of sleep spindles reflects a frequency decrease or a change in the relative predominance of slow over fast sleep spindles (Aeschbach et al., 1997; Wei et al., 1999). Moreover, there is no direct evidence for the detectability of the above described frequency evolution of sleep spindles during habitual (non-displaced) sleep periods. A study examining the overnight evolution of sleep spindle frequencies, but without differentiating slow and fast sleep spindle events, revealed a sleep time-dependent increase in frequency (Himanen, Virkkala, Huhtala, & Hasan, 2002). This pattern evidently contrasts the convergent findings reported by studies implementing chronobiology protocols and systematic sleep displacement. The discrepancy might reflect a contamination of two different factors: change in sleep spindle frequencies and relative slow over fast sleep spindle incidence during the course of the night.

Our aim is to test the feasibility of measuring the circadian modulation of sleep spindle frequencies in one-night-records of habitually timed sleep periods in different age groups by analysing its overnight dynamics (change over consecutive sleep cycles) and by comparing night-time sleep spindle frequency with afternoon nap sleep spindle frequency. We also aim to provide a differential analysis of slow and fast sleep spindles and depict the potential difference between oscillatory deceleration of slow and/or fast sleep spindles and alternatively, the change in the relative predominance of slow over fast sleep spindle events. In order to achieve these goals, we use an already established procedure of individualizing slow and fast sleep spindle frequencies with high frequency resolution (Bódizs et al., 2009). Given the already evidenced circadian modulation and the reported melatonin- and/or temperature-dependence of sleep spindle frequency, we hypothesize that:

1. Overnight dynamics in sleep spindle frequency is characterized by a U-shaped distribution (sleep spindles are slower in the middle of the habitual sleep period, as compared to the first and the last sleep cycles)
2. Middle night slowing of sleep spindle frequency is reduced in aged subjects as compared to young participants
3. Estimated phase of the nadir in sleep spindle frequency is delayed in teenagers and young adults, as compared to children and middle aged adults
4. Night sleep spindles are slower than nap sleep spindles.

## 2 METHODS

### 2.1 Subjects and databases

Multiple, published databases are used in this study. The Munich-Budapest database of sleep records consists of 251 night-timed polysomnography registrations (Bódizs et al., 2017). Subjects’ age varies between 4 and 69 years (122 females, mean age: 25.73 years). Participants of the night sleep record dataset slept at their habitual sleeping time in the laboratory (N = 208) or in their homes (recorded by ambulatory polysomnography, N = 43) on two consecutive nights. In order to attenuate the first-night effect (Agnew, Webb, Williams, & Miller, 1966), only the second night data were used. Caffeine containing, but not alcoholic beverages were allowed in the morning hours in our adult subjects (maximum 2 cups of coffee/subjects before noon), whereas some of the participants who were light smokers (N = 8) were allowed to smoke during the day. The nap sleep records (N = 112) stem from studies on the effects of napping on memory consolidation (Genzel et al., 2014, 2012), as well as from a study analysing the relationship between nap sleep spindles and intelligence (Ujma et al., 2015), but only baseline and not post-learning records are analysed in the present investigation. Age of the napping subjects varied between 18 and 30 years (mean: 23.72 years). Women involved in the nap sleep studies (N = 30) were recorded twice: afternoon naps took place during the first and the third week of their menstrual cycle (early-follicular and mid-luteal phases, respectively with hormonal blood tests assuring this assumption). Data of the two nap records of these subjects were averaged before statistical analyses. All subjects were healthy and free of any medications, except contraceptives in some of the women in the reproductive age. Details of the recording procedures are reported in Table 1. All subjects (night sleepers and nap sleepers) were free of acute and chronic medical conditions, including the history of sleep disorders. The research protocols were approved by the Ethical Committee of the Semmelweis University (Budapest, Hungary) or the Medical Faculty of the Ludwig Maximilians University (Munich, Germany) in accordance with the Declaration of Helsinki. All subjects or the parents of the underage participants signed informed consent for the participation in the studies.

**Table 1.** Details of the recording procedures of different subsamples

| Nap/<br>night | Subsample | Original setting/aim | Nr. of<br>subjects<br>(females) | Age<br>range<br>(ys) | Available EEG<br>derivations (10-20 system) | Recording<br>apparatus | Precision<br>(bit) | Hardware prefiltering (Hz) | Sampling<br>rate<br>(Hz/channel) | Recording<br>software |
| --- | --- | --- | --- | --- | --- | --- | --- | --- | --- | --- |
| Night sleep | MPIP<sup>1</sup> – I | Lab/sleep & IQ | 95 (43) | 18–69 | Fp1, Fp2, Fpz, AF1, AF2, F3, F4, Fz, F7, F8, C3, C4, Cz, P3, P4, Pz, T3, T4, T5, T6, O1, O2 | Comlab 32 Digital SleepLab | 8 | 0.53–70 | 250 | Brainlab 3.3 |
|  | MPIP<sup>1</sup> – II |  | 20 (12) |  | Fp1, Fp2, F3, F4, C3, C4, P3, P4, O1, O2 |  |  |  |  |  |
|  | PPCU<sup>2</sup> – I | Home/Williams syndrome study (controls included here) | 20 (14) | 6–28 | Fp1, Fp2, Fpz, F3, F4, F7, F8, Fz, C3, C4, Cz, T3, T4, T5, T6, P3, P4, Pz, O1, O2, Oz | SD-LTM 32BS (Micromed Ltd, Italy) | 22 | 0.15–250 (plus <463.3 Hz digital antialiasing filtering before downsampling from 4096 to 1024 Hz) | 1024 | BRAIN QUICK System PLUS (Micromed) |
|  | PPCU<sup>2</sup> – II | Home/Adolescent sleep | 23 (12) | 15–22 |  |  |  |  |  |  |
|  | SU<sup>3</sup> – I | Lab/sleep & IQ, sleep spindle methodology, wake-sleep transition analysis | 49 (19) | 17–55 | Fp1, Fp2, F3, F4, F7, F8, Fz, C3, C4, Cz, T3, T4, T5, T6, P3, P4, O1, O2 | Flat Style SLEEP La Mont Headbox, HBX32-SLP preamplifier (La Mont Medical) | 12 | 0.5–70 | 249 | Datalab (Medcare) |
|  | SU<sup>3</sup> – II | Lab/nightmare study (controls included here) | 16 (7) | 19–21 | Fp1, Fp2, F3, F4, Fz, F7, F8, C3, C4, Cz, P3, P4, Pz, T3, T4, T5, T6, O1, O2 | Brain-Quick BQ132S (Micromed) | 12 | 0.33–1500 (plus <450 Hz anti-aliasing digital filtering before downsampling from 4096 to 1024 Hz) | 1024 | System 98 (Micromed) |
|  | SU<sup>3</sup> – III | Lab/home/ children’s dreaming | 29 (15) | 3.84–8.42 |  | Brain-Quick BQ132S/ SD LTM 32BS (Micromed) | 12/22 | 0.33–1500/0.15–250 (plus <450/<463.3 Hz anti-aliasing digital filtering before downsampling from 4096 to 1024 Hz) |  | System 98/System Plus Evolution (Micromed) |
| Nap sleep | MPIP – III | Lab/Nap sleep & memory consolidation | 55 (35) | 18–30 | (EOG) C3, C4 | Comlab 32 Digital SleepLab | 8 | 0.53–70 | 250 | Brainlab 3.3 |
|  | MPIP – IV | Lab/Nap sleep & IQ | 79 (0) | 18–30 | F3, F4, C3, C4, O1, O2 | Comlab 32 Digital SleepLab | 8 | 0.53–70 | 250 | Brainlab 3.3 |

<sup>1</sup>Max Planck Institute of Psychiatry, Munich, Germany; <sup>2</sup>Pázmány Péter Catholic University, Budapest Hungary; <sup>3</sup>Semmelweis University Budapest, Hungary;
Continuous underlined symbols: frontal recording sites (missing frontal EEG derivations in the nap database MPIP-III were substituted with the EOG channel in order to define slow spindle ranges); Dotted underlined/grey-shadowed symbols: frontal and centroparietal recording sites, respectively

### 2.2 EEG processing

Records were scored according to standard criteria of sleep-wake states (Berry et al., 2018), followed by artefact-removal on a 4 seconds basis. Sleep cycle segmentation was based on reported criteria (Aeschbach & Borbély, 1993). EEG signals (mathematically-linked mastoid reference) of the non-artefactual N2 and N3 sleep periods were subjected to the Individual Adjustment Method (IAM) of sleep spindle analysis (Bódizs et al., 2009) in chunks based on successive sleep cycles of night-time records and as a whole in case of nap sleep. Only the frequency outputs of the IAM is reported in the present study.

The IAM consists of a high-resolution (1/16 Hz) frequency adjustment of characteristic slow- and fast sleep spindle frequencies, based on the zero-crossing points of the averaged second order derivatives of the NREM sleep EEG amplitude spectra (magnitude of the 4 s, Hanning tapered mixed-radix Fast Fourier Transforms). That is, amplitude spectra of NREM sleep EEG with 0.25 Hz frequency resolution were subjected to a numerical derivation procedure as follows: a second-degree polynomial curve fitting was performed using all sets of successive bin triplets (0.75 Hz), with an overlap of 2 bins (0.5 Hz) in the 9–16 Hz range. Derivatives were calculated in the middle of the triplets by using the rules of derivation for quadratic formulas, returning practically the slopes of the curves fitted to the triplets. This procedure is repeated on the series composed by the first derivatives in order to obtain the second derivatives of the amplitude spectra. Second derivatives were averaged over the EEG recording locations resulting in one averaged second derivative function per subject and sleep cycle (NREM period). The zero crossing points of these averaged second derivatives were refined on the frequency scale of the high-resolution spectra (1/16 Hz) and considered as lower and upper frequency boundaries of slow and fast sleep spindles if the following criteria were fulfilled. Fast sleep spindle spectral peaks were defined if

a. there was a negative peak in the averaged second derivative (positive peaks are reverted and negative in the second order derivatives) at the corresponding frequencies
b. the peak was of highest frequency (there is no spectral peak with higher frequency in the spindle range)
c. the amplitude spectra at the assumed peak frequencies was higher in centroparietal as compared to frontal recording locations (Fig. 1).

**FIGURE 1.**
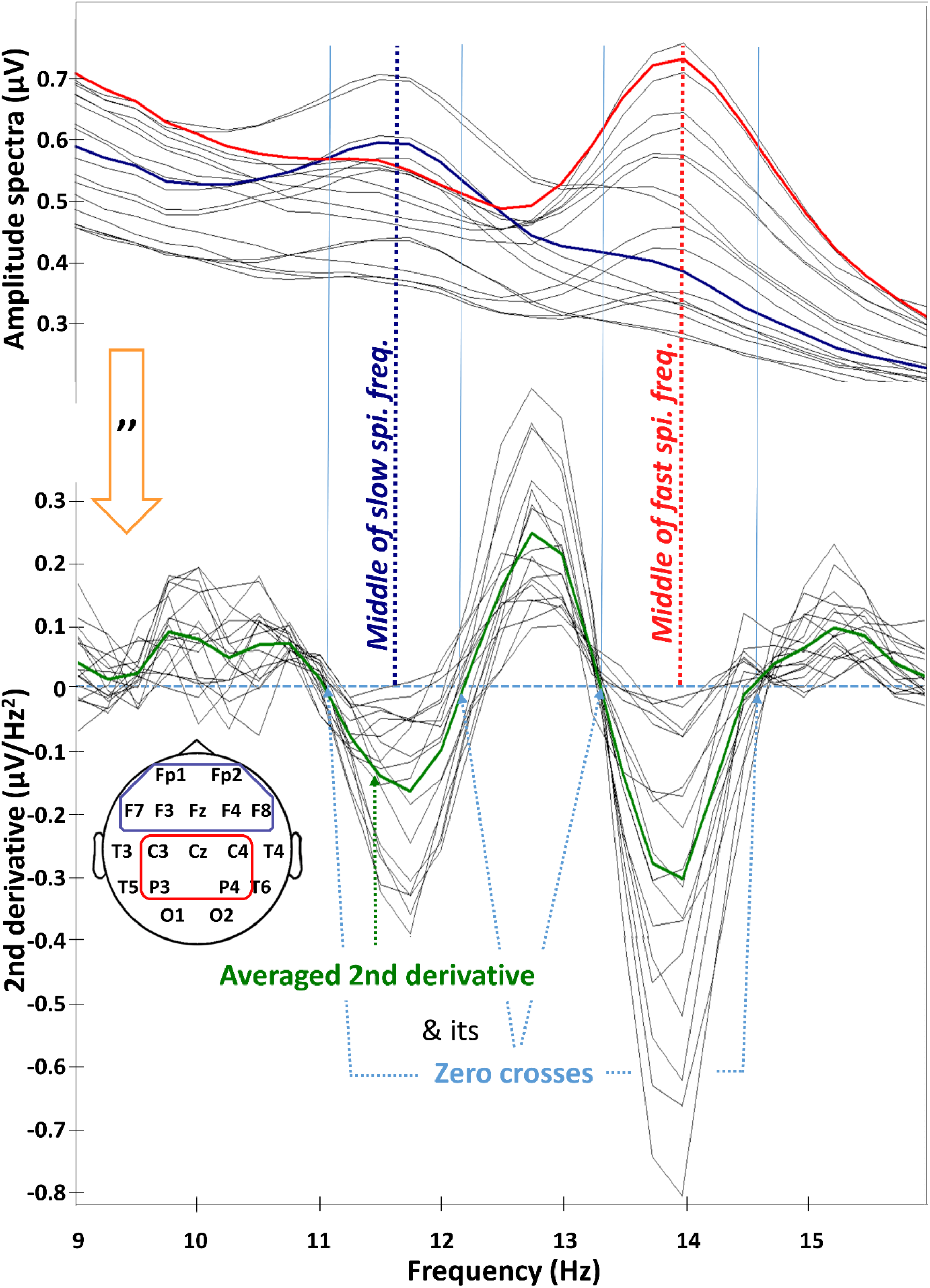
Critical steps in determining the individual slow and fast sleep spindle frequencies. Amplitude spectra, as well as their frontal and centroparietal averages (continuous blue and red lines, respectively) are calculated by means of (mixed-radix) Fast Fourier Transformation (4 sec, Hanning window, 2 sec overlap, see upper panel). The definition of frontal and centroparietal regions is provided in the lower insert (circle representing the head model). Amplitude spectra are submitted to a numerical derivation procedure in two steps, resulting in second derivatives (”, lower panel). The latter are averaged (continuous green line) followed by the detection of their zero crosses (interpolated on a scale of 1/16 Hz, see dotted, light blue arrows). Last, the individual spectral peak frequencies are determined by combining the frontal and parietal spectral peaks with the zero crosses determined above. Note that the procedure is the first step of the Individual Adjustment Method (IAM) of sleep spindle analysis.

Frontal recording locations were defined as all available scalp contacts whose symbol begins with F or AF. In turn, centroparietal region was defined by all available contacts labelled with C or P. Fast sleep spindle frequencies were unambiguously detected in all subjects and sleep cycles.

In contrast, slow sleep spindle spectral peaks were defined if

a. there was a negative peak in the averaged second derivative at the corresponding frequencies
b. the peak was of lower frequency (there was at least one peak with higher frequency in the spindle range)
c. the relative difference in centroparietal and frontal amplitude spectra at the assumed peak frequencies was (relatively) lower as compared to this difference seen at the fast sleep spindle frequencies.

In cases in which more than two peaks were present, we based our selection on frontal and centroparietal dominances of the respective frequencies, relying on the amplitude spectra. That is, the peak with most pronounced frontal and centroparietal dominance in amplitude spectra, were considered as slow and fast sleep spindle frequencies, respectively. Lack of a definitive slow spindle peak was handled by searching the relative maxima of the frontal minus centroparietal amplitude spectral values, the slow spindle spectral peaks being defined at the edges of the associated bulge of second-order derivatives (which latter approach but do not reach zero in these cases; see Suppl. fig. 1) (Ujma et al., 2016). Last, but not least the spectral peaks of slow and fast sleep spindles could partially overlap in some cases. In such cases the intersection of slow and fast sleep spindle-related spectral peaks was considered as a border frequency.

Slow and fast sleep spindle frequencies were defined as the middle of the respective frequency bands (arithmetic mean of lower and upper frequency boundaries).

The estimated phases of the nadirs of slow and fast sleep spindle frequencies was assessed as follows:

i. finding the sleep cycle characterized by the individual minima in sleep spindle frequency
ii. determining the local time of day of the middle of the respective sleep cycle (NREM-REM period) by recovering the start time of the respective sleep recording

transforming time of day values (HH:MM:SS) to hours and fractions of hours relative to midnight (that is times of day before midnight became negative, whereas those after midnight emerged as positive values; higher values indicate later phases of the nadir in sleep spindle frequency).

We could recover the start times of the original recording in N = 248 subjects (98.8% of the subjects). In order to provide a composite measure of the assumed circadian phase of sleep spindle frequency we calculated the individual mean of the phases of slow and fast sleep spindle frequency minima.

Slow wave activity (SWA) was defined as the power spectral density of 0.75–4.5 Hz EEG activity (sum of the bin power values) by using a (Mixed-radix) Fast Fourier Transformation routine on artefact-free, 4 sec, Hanning-tapered windows (2 sec overlap). Analysis was performed on the left frontal EEG recording location (F3), and averaged in chunks of NREM periods of complete sleep cycles.

### 2.3 Statistical analyses

Subjects of the night sleep study were classified according to the following age ranges (Bódizs et al., 2017): children (4 years ≤ age < 10 years; N = 31, 15 females), teenagers (10 years ≤ age < 20 years; N = 36, 18 females), young adults (20 years ≤ age < 40 years; N = 150, 75 females) and middle-aged adults (40 years ≤ age ≤ 69 years; N = 34, 14 females). The sleep cycle effect in oscillatory spindle frequency was tested with the control of the between-subject factors sex and age group, the within-subject factor spindle type (slow/fast), as well as all possible interactions of the above mentioned predictors, by using the general linear model (GLM) approach. Associations between sleep spindle deceleration and age, as well as between bedtime and NSSF were tested by the Pearson product-moment correlation coefficients. The comparability of nap and night sleep records was ensured by matching a subset of the night sleep records (the young adult group, N = 141, 69 females) to the nap sleep database (N = 108, 29 females) in terms of age and there were no significant difference between the two groups (t(247) = −0.318, p =.75). Besides our main focus (the nap vs night between subject factor), sex (between-subject), and spindle type (slow/fast, within-subject) were included in the GLM. In spite of the homogenous age of the subjects of the nap vs night sleep study, the age factor was included in an additional statistical model as a continuous predictor.

## 3 RESULTS

### 3.1 Overnight dynamics in sleep spindle frequency

Sleep architecture data indicated typical sleep composition (Suppl. table 1). Number of complete sleep (NREM-REM) cycles in night sleep records varied between 2 and 5 (mean: 4.24). 251 subjects had at least 2, 249 had minimum 3, 239 had ≥ 4, and 75 had 5 complete sleep cycles. In order to keep the number of subjects high, first we focused on NREM periods of the first four sleep cycles (~6.8 hours after sleep onset, N = 239). Mean (standard deviation) duration of sleep cycles was 102.22 (36.64), 106.33 (30.71), 103.97 (23.86), 95.68 (26.94), and 81.59 (21.13) minutes for cycle 1, 2, 3, 4, and 5 respectively. No significant age, sex, cycle main effects or interactions were revealed for sleep cycle durations (GLM for cycles 1–4, repeated measures with the categorical predictors age group and sex).

Significant sleep cycle (cycles 1–4, N = 239) effects (F = 18.57; d.f. = 3, 693; p < .001) revealed a U-shaped dynamic in spindle frequencies, with highest values in the first and the fourth cycles and lowest values in the second and third cycles. This pattern was evident for both slow and fast sleep spindle types, characterized by overall frequency drops of 0.09 and 0.13 Hz, respectively (Fig. 2; post-hoc Fisher LSD tests indicate decelerated sleep spindles in cycles 2 and 3, as compared to cycles 1 and 4 in terms of both slow and fast spindles, whereas cycles 1 and 4 do not differ significantly). The sleep cycle effect interacted with the between-subject factor age group (F = 2.07; d.f. = 9, 693; p = .03), indicating an attenuated spindle deceleration in middle aged adults (flattening of the U-shaped dynamic), but also in the lack of this pattern in the time course of slow sleep spindles in girls below 10 years of age (Fig. 3). The age-related flattening of overnight dynamic is supported by the negative correlation between age and a measure of deceleration of spindles (mean spindle frequency in cycles 1 and 4 minus mean spindle frequency in cycles 2–3): r = −.12 (p = .049) for slow and r = −.21 (p = .001) for fast types. Nominal sleep spindle deceleration reached 0.18 and 0.20 in case of slow and fast sleep spindles of teenagers, respectively, whereas it was only 0.03 and 0.04 in middle aged adults.

**FIGURE 2.**
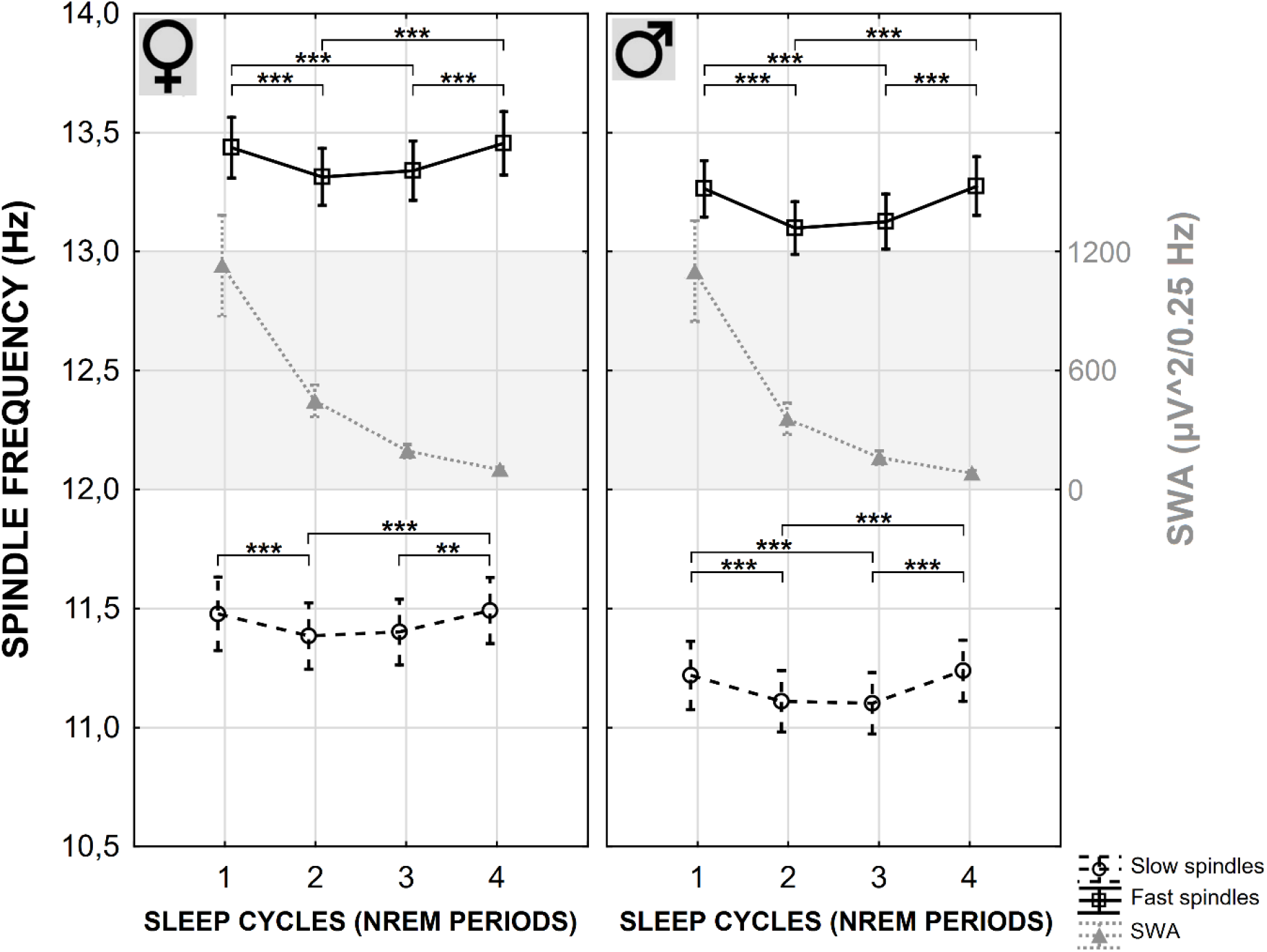
The U-shaped dynamics in the overnight evolution of sleep spindle frequencies. Slow and fast sleep spindles (broken and continuous black lines, respectively) decelerate in sleep cycles 2 and 3, whereas cycles 1 and 4 are characterized by higher values. This effect is evident for both females (♀) and males (♂), despite the overall higher oscillatory frequencies of sleep spindles in females. The U-shaped overnight dynamics of the oscillatory frequency of sleep spindles in successive sleep cycles, clearly differs from the exponential decay of slow wave activity (SWA, shaded area, grey, dotted lines). The latter is known as the most reliable indicator of sleep homeostasis.

**FIGURE 3.**
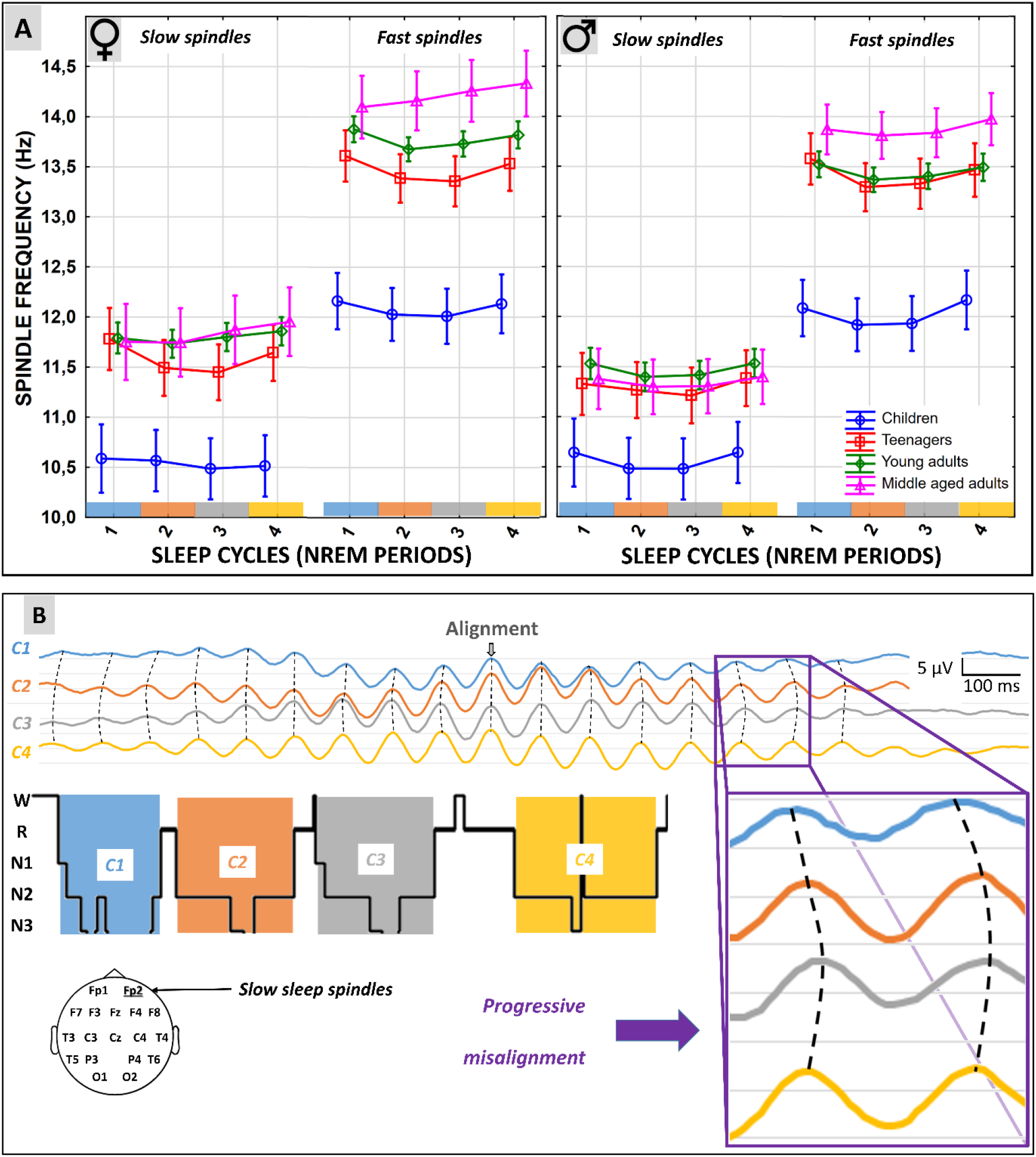
Overnight dynamics of slow and fast sleep spindle frequencies as functions of age, and sex. A. The U-shaped overnight dynamics characterized by the highest oscillatory frequencies in the first and in the fourth sleep cycles is seen in all instances, except slow sleep spindles in girls below 10 years of age. Sleep spindles become faster with increasing ages, whereas the mid-sleep deceleration is attenuated in middle aged participants. B. Example for the deceleration of sleep spindles in the middle of the habitual sleep period in a 20 years old male. Slow sleep spindles measured at recording location Fp2 are averaged to the maxima of the spindle-filtered signal. Averaged sleep spindles of successive sleep cycles are aligned according to their maxima. Note the progressive misalignment of averaged sleep spindle waves of sleep cycles 2 and 3, indicating their decelerated frequency.

In sharp contrast with the U-shaped dynamics of sleep spindle frequencies, SWA was characterized by a continuous, exponential decline (Cycle effect: F = 332.42; d.f. = 3, 666; p < .001). Earlier sleep cycles were characterized by significantly higher SWA as compared to later ones, except cycles 3 and 4, which did not differ significantly (post-hoc Fisher LSD tests).

As expected, sleep spindle frequency varied between age groups (F = 66.46; d.f. = 3, 693; p < .001) and as a function of sex (F = 8.94; d.f. = 1, 693; p = .003), indicating increasing oscillatory frequencies with increasing ages, and higher frequencies in females as compared to males, respectively (Fig. 3).

A subset of records contained a complete 5^th^ sleep cycle (N = 75, that is 29 children, 23 teenagers, 22 young adults, and 1 middle aged adult). The above statistical analyses were extended over the 5^th^ sleep cycle by relying on this subset of records, with the evident exclusion of the single middle aged subject (N = 74). Results still reflected a significant sleep cycle effect on spindle frequency (F = 17.08; d.f. = 4, 272; p < .001), but not interaction between cycle effect and age group (because the middle aged group of subjects with flattened U-shaped dynamic was not included in this latter analysis: F = 0.50; d.f. = 8, 272; p = .85). We implemented the analysis on this subset of records in order to focus on the 5^th^ sleep cycle. Results revealed that sleep spindle frequencies measured in this latter sleep cycle exceeded all other sleep spindle frequencies measured in cycles 1–4, according to post-hoc Fisher LSD tests.

In order to provide a further test of our assumptions we performed an additional analysis of across-night effects in sleep spindle frequencies for individually determined first, middle and last sleep cycles on the whole sample (N = 251), which lead to the same results as reported above: lower frequencies in the middle of the sleep period and an interaction of cycle position effects with the between subject factor age group (Suppl. fig. 2; Suppl. table 2).

### 3.2 The phase of the nadir in sleep spindle frequency (NSSF)

The sample mean of the phase of the NSSF was 2.63 (SD = 2.16) and 2.49 (SD = 2.01) hours for slow and fast sleep spindles, respectively. That is most of the participants reach their lowest sleep spindle frequencies roughly two and a half hours after midnight (the sample mean of the individually averaged phases of slow and fast NSSF was 2.56 hours). A GLM targeting the effects of age group, sex, spindle type and their interaction on NSSF resulted in a significant main effect of age group (F = 5.30; d.f. = 3, 240; p = .001). Based on the means and standard errors depicted in Figure 4, it is evident, that the hypothesized age effects are best approximated by slow sleep spindle frequencies: phase of the NSSF is seen circa one hour after midnight in children, whereas it reaches 3 hours in teenagers, followed by a slow decrease to 2.5 hours in upcoming age groups. Post-hoc Fisher LSD tests revealed a significantly earlier phase of the NSSF (slow spindles) in children as compared to all other age groups. Nominal differences among teenagers and adults (young and middle aged) did not result in significant post-hoc effects.

**FIGURE 4.**
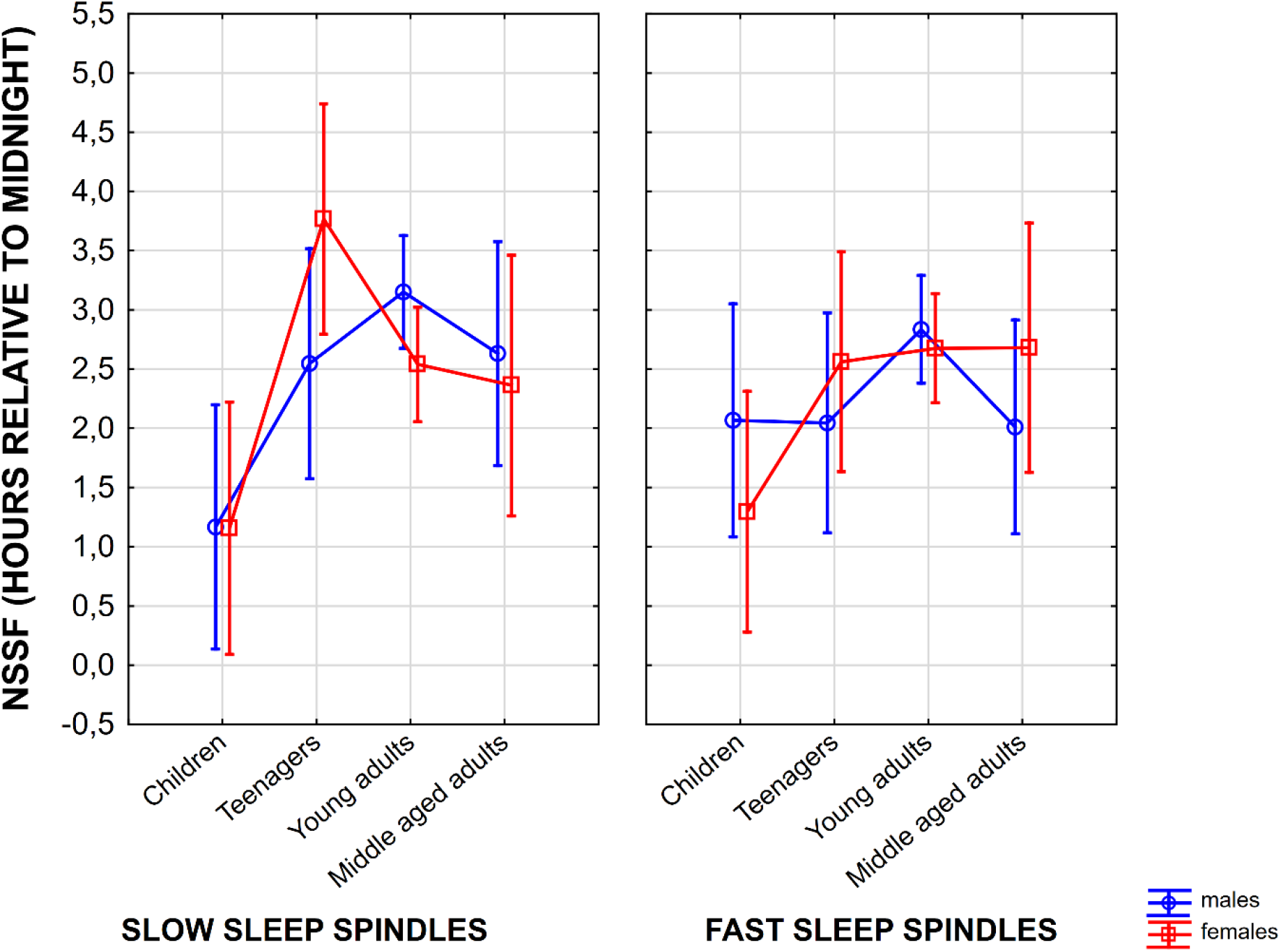
Phase of the nadir in sleep spindle frequency (NSSF) as expressed in fraction of hours relative to midnight. Vertical bars denote 95% confidence intervals.

In order to test the potential dependence of NSSF on bedtimes, which would clearly confuse the interpretation of our findings, a Pearson correlation coefficient was calculated. The latter indicated no relationship between bedtime and NSSF (r = .02; p = .68), suggesting that later bedtimes do not lead to later NSSF.

### 3.3 Nap vs night sleep spindle frequency

Nap sleep architecture is given in Supplementary table 3. Nap sleep spindles were faster than night sleep spindles (F = 72.2; d.f. = 1, 249; p < .001). Although this effect is evident for both spindle types (Fig. 5), day vs night slow spindle frequency differences are higher than corresponding fast spindle differences (0.57 vs 0.39 Hz; interaction of nap vs night and spindle type: F = 4.6; d.f. = 1, 249; p = .033). Oscillatory frequency of sleep spindles was higher in females as compared to males (F = 22.7; d.f. = 1, 249; p < .001), but this sex-difference did not interact with the nap vs night effect, which is the focus of the current study. We obtained essentially the same results after including age as a continuous predictor in the model. Nap sleep spindle frequencies exceeded night sleep spindle frequencies (F = 70.14; d.f. = 1, 244; p < .001), especially, but not exclusively in case of slow sleep spindles (F = 5.01; d.f. = 1, 244; p = .025). Faster sleep spindles were recorded in women as compared to men (F = 20.29; d.f. = 1, 244; p < .001). Age did not predict sleep spindle frequencies in this age range (F = 1.08; d.f. = 1, 244; p = .29). Given the fact that this latter analyses involved a between-subject comparisons of records obtained with different hardwares, we performed an additional statistical test by using a subsample of the night sleep EEG registrations performed with the same recording system as the one used in nap sleep studies. This latter analysis fully replicated the results reported under this subheading (Supplementary table 4).

**FIGURE 5.**
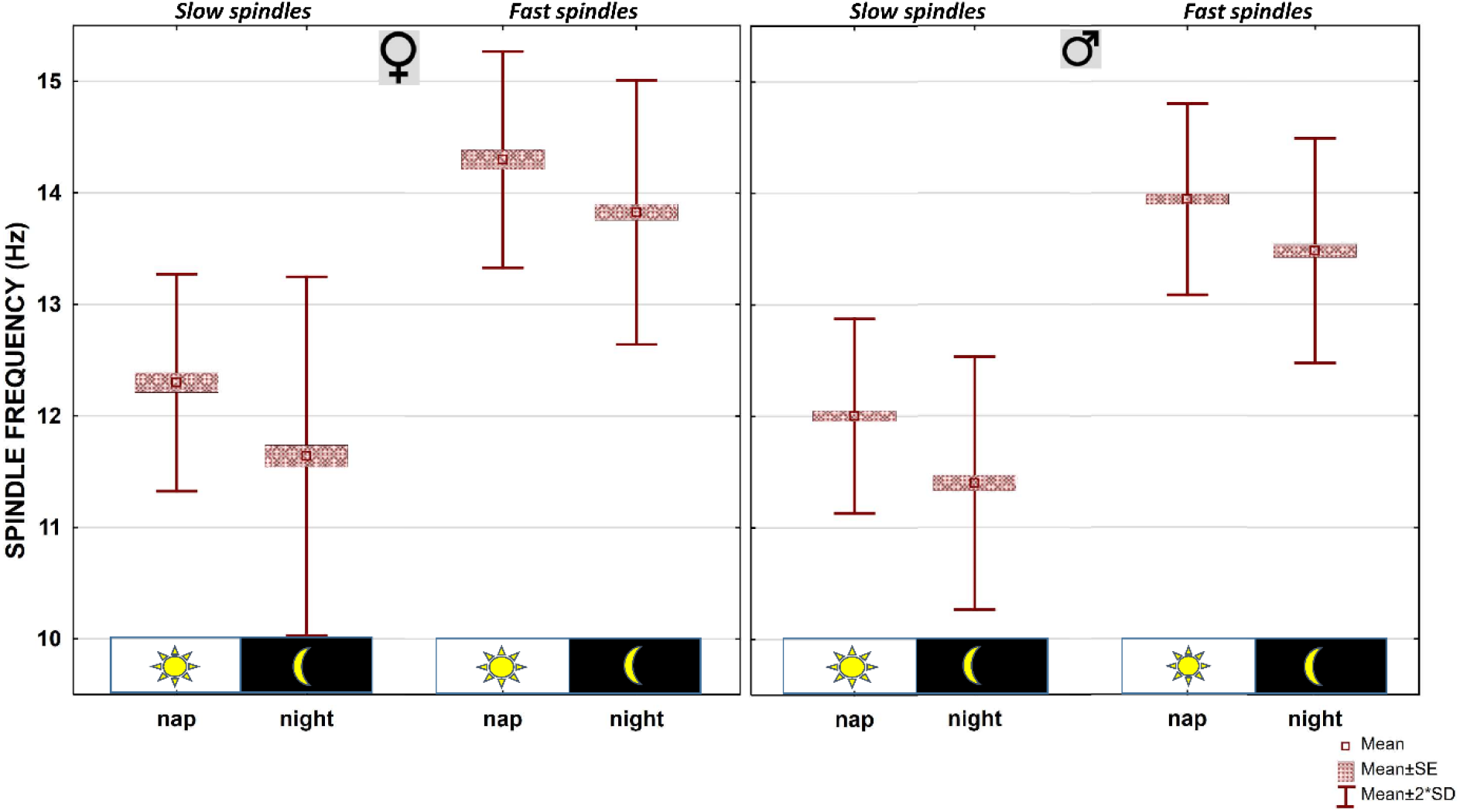
Nap sleep spindles are higher than night sleep spindles. Individual-specific slow- and fast sleep spindles are compared in two groups of age- and sex-matched subjects sleeping during the afternoon and during the night, respectively.

## 4 DISCUSSION

Our findings indicate that sleep spindles decelerate in the middle of the habitual sleeping period, by roughly 0.1 Hz. This effect is evident for both slow and fast sleep spindles, thus it is not caused by the redistribution of slow and fast sleep spindle occurrences during the night. Younger subjects are characterized by more accentuated deceleration of sleep spindling in the middle of their sleeping period (the nominal values of spindle deceleration in the middle of the night sleep period of teenagers are around 0.2 Hz). This U-shaped dynamics, characterized by an equality of the first and the fourth sleep cycle, and the decrease in the second and the third cycle cannot be explained by homeostatic sleep regulatory mechanisms, which latter would imply a strictly decreasing or increasing function. Findings reported by the relevant forced desynchrony (Wei et al., 1999), constant routine (Knoblauch et al., 2005) and sleep displacement (Aeschbach et al., 1997) studies suggest the involvement of circadian regulation in modulating sleep spindle frequency in humans. Former studies analysing the overnight dynamics of sleep spindle frequency in habitually-timed polysomnography records, did not deliberately discern slow and fast sleep spindles (Himanen et al., 2002), thus could at least partially reflect the changing dominance of slow over fast sleep spindles. The latter was reported in former studies, suggesting that both the incidence and the amplitude of slow spindles decreased over successive sleep cycles, whereas an opposite trend was observable for fast sleep spindles (Bódizs et al., 2009; Purcell et al., 2017). Overnight changes in slow and fast sleep spindle incidence might explain the roughly 2 Hz acceleration of sleep spindles reported by Himanen et al. (2002).

Frontal and centroparietal regions are the main sources of slow and fast sleep spindles, respectively. This topographical feature was used as a priori criteria of defining slow and fast sleep spindle frequencies in our current paper (Fig. 1), but not investigated as a dependent variable. That is, we assumed that a slow or a fast sleep spindle frequency is individual-, but not region-specific. Given the fact that topographical differences in oscillatory frequencies within the individual-specific slow or fast sleep spindle domain cannot be ruled out, this issue merits further attention in later studies.

The deceleration of slow and fast sleep spindles we report in our current study could be a result of the increased melatonin levels and the associated decrease in core body temperature in the middle of the habitual sleep period. The acrophase of the human plasma melatonin rhythm is known to emerge between 2:00 and 5:00 AM (Voultsios, Kennaway, & Dawson, 1997). In addition, melatonin receptors are expressed in the reticular thalamic nucleus (Ng, Leong, Liang, & Paxinos, 2017), that is a critical neuroanatomical structure in the process of sleep spindle generation (Fan, Liao, & Wang, 2017). Thus, both the timing and one of the target organs of melatonin release are ideally suited to support the involvement of the pineal hormone in shaping the evolution of sleep spindle frequency during the night. Last, but not least, sleep spindle frequency varies as a function of locally manipulated brain temperature (Csernai et al., 2019), whereas melatonin is known for its hypothermic effect (Marrin, Drust, Gregson, & Atkinson, 2013). As a consequence, the hypothermia induced by increased melatonin levels in the middle of the sleeping period is potentially involved in the deceleration process of sleep spindling revealed in the current report. This assumption is further supported by differences in sleep spindle frequency activity during and outside melatonin secretory phase (Knoblauch et al., 2005), as well as by the findings indicating that spindle frequency reaches its nadir at the trough of the body temperature cycle (Wei et al., 1999).

The age-dependent attenuation of sleep spindle deceleration in the middle of the habitual sleep period reported herein is reminiscent of the age-related decline in the amplitude of the circadian rhythm (Hood & Amir, 2017), the flattening of the body temperature rhythm in the aged (Weitzman, Moline, Czeisler, & Zimmerman, 1982), as well as the associated decline in melatonin release (Waldhauser et al., 1988). Decreased circadian modulation and melatonin production might fail to induce an efficient reduction in core body temperature and consequently a suboptimal modulation of sleep spindle frequency. Our findings cohere with the forced desynchrony study reporting that older subjects express notably smaller circadian variation of sleep spindle frequency than young subjects (Wei et al., 1999). However, the reason for a lack of the above detailed U-shaped distribution in the frequency of slow sleep spindles in young girls do not cohere with the above reasoning.

Sleep spindles speed up in the fourth sleep cycle and continue to accelerate on the rising limb of the circadian cycle up to the 5^th^ sleep cycle in the subset of subjects having five complete NREM-REM periods in our dataset. Thus, we assume that sleep spindles are even faster during daytime as compared to night-time sleep. This coheres with our findings reported in the nap study. That is, in addition to the reported deceleration of spindle oscillations in the middle of the habitual night sleep period, we report accelerated spindle frequency during afternoon naps as compared to nocturnal averages (cycles 1–4). The finding that the oscillatory frequency of nap sleep spindles exceeds that of night spindles further strengthens our assumption that the above discussed sleep cycle effects might reflect circadian modulation and/or melatonin release. Core body temperature is known to be higher during afternoon hours as compared to late night periods (Baehr, Revelle, & Eastman, 2000), whereas melatonin release is evidently at its lowest level during these daytime hours (Aulinas, 2000). Thus, the acceleration of sleep spindles in afternoon naps coheres with our assumption regarding the circadian regulation of the duration of thalamocortical hyperpolarization-rebound sequences. To the best of our knowledge this is the first study explicitly reporting a nap vs night sleep difference in spindle frequencies.

We hypothesized that the phase of the middle night drop in sleep spindle frequency could serve as an EEG-marker of the circadian phase in humans. Available data in the current set is only suitable for an indirect test of this assumption. That is, NSSFs were found to be independent from bedtimes, which might suggest a time-of-day rather than a sleep-dependent effect. Moreover, we tested the known age-effects in circadian phase by comparing the different age groups. Our findings partially support the hypothesis: children were characterized by earlier phases than teenagers or adults. Although, this latter finding coheres with known age-differences in circadian phase, additional age effects, namely the phase advancement in middle aged adults was not unequivocally supported by our findings. This could reflect the lack of aged subjects in our sample or the insufficient precision of measuring the phase of the nadir of sleep spindle frequency (in the middle of the sleep cycle characterized by lowest sleep spindle frequency). Later studies could invoke a more instantaneous phase measure, providing investigators with increased temporal resolution. Given the partial statistical support, we consider our measure as a potentially suitable EEG-index of circadian phase, which might bridge the methodological and conceptual gap between chronobiology and somnology in future investigations.

Besides of the U-shaped overnight dynamics and afternoon nap effect, formerly reported sex-differences and age-effects in sleep spindle frequencies were also supported by our current analyses. That is, sleep spindles were of higher frequency in females as compared to males, an effect which was already reported by different research groups (Markovic, Kaess, & Tarokh, 2020; Ujma et al., 2014). Moreover, sleep spindle frequencies were significantly lower in prepubertal ages (below 10 years), as compared to teenagers and adults. This effect was also reported by our own former study (Ujma et al., 2016), as well as by other teams analysing sleep spindles in children (Campbell & Feinberg, 2016). The convergence of these findings with the available reports in the literature strengthens the validity of our current approach and provides further empirical support for the recent hypothesis on the neurodevelopmental relevance of sleep spindle frequency(Zhang et al., 2021).

Given the fact that individual-specific slow and fast sleep spindle bands are roughly 1 Hz wide each (Bódizs et al., 2009), the herein reported ~0.1 Hz overnight changes in frequency do not compromise the trait-like stability of night sleep spindle measures. However, the ~0.5 Hz acceleration of sleep spindles during daytime naps as compared to night sleep needs indeed further attention from the perspective of trait-like stability.

The strength of our study is the high number of subjects and wide age range, whereas limitations are the lack of repeated day vs night sleep measurements in the same groups of subjects, the lack of controlling several variables related to menstrual cycles (cycle phase, contraceptive use), as well as the lack of valid circadian measures (body temperature, melatonin release) against which we could perform tests of convergent validity. Although some of the subsamples we use are of lower sampling rate as compared to the current standards of the American Academy of Sleep Medicine (500 Hz), herewith we only focus on relatively lower frequency oscillations (at least 15 times lower as compared to our lowest sampling rate), which (together with the antialising hardware filters) provide sufficient technical support for our conclusions. We consider our findings as a first step in defining an EEG-index of the circadian modulation of sleep, which could efficiently complete the already available and widely used measure of sleep homeostasis. We hypothesize that the reliable measurement of the overnight dynamics of slow and fast sleep spindle frequencies might convey information about the amplitude and perhaps the phase of the circadian rhythm in future translational and clinical studies, including investigations on patients suffering from circadian rhythm sleep disorders.

## ACKNOWLEDGEMENTS

Research supported by the Hungarian National Research, Development and Innovation Office (K-128117; https://nkfih.gov.hu/about-the-office), the Higher Education Institutional Excellence Program of the Ministry of Human Capacities in Hungary, within the framework of the Neurology thematic program of the Semmelweis University (http://semmelweis.hu/english/), the Netherlands Organization for Scientific Research (NWO; https://www.nwo.nl/en), the European Cooperation in Science and Technology (COST Action CA18106; https://www.cost.eu/), as well as the general budgets of the Institute of Behavioural Sciences, Semmelweis University (http://semmelweis.hu/magtud/en/) and the Max Planck Institute of Psychiatry (https://www.psych.mpg.de/en). The funders had no role in study design, data collection and analysis, decision to publish, or preparation of the manuscript. Authors wish to thank Dorina Ali and Dávid Rottmayer for their assistance in data analysis.

## CONFLICT OF INTERESTS

No conflicts of interest declared.

## AUTHOR CONTRIBUTIONS

R. B. conceived the study; R. B., P. S., I. K., L. G., and M. D. contributed to data collection; R. B., C. G. H., P. P. U., P. S., and F. G. contributed to data analysis; all authors drafted the manuscript, critically revised the major intellectual content and approved the final version of the paper.

## Notes

### Competing Interest Statement

The authors have declared no competing interest.

