## Supplemental Data 1 for "Sleep spindle frequency: overnight dynamics, afternoon nap effects, and possible circadian modulation"

**Supporting information for the paper entitled "Sleep spindle frequency: overnight dynamics, afternoon nap effects, and assumed circadian modulation"**

Róbert Bódizs[1,2], Csenge G. Horváth[1], Orsolya Szalárdy[1,3], Péter P. Ujma[1,2], Péter Simor[1,4,5], Ferenc Gombos[6,7], Ilona Kovács[6], Lisa Genzel[8], Martin Dresler[8]

*[1]Institute of Behavioural Sciences, Semmelweis University, Budapest, Hungary*

*[2]National Institute of Clinical Neurosciences, Budapest, Hungary*

*[3]Institute of Cognitive Neuroscience and Psychology, Research Centre for Natural Sciences, Budapest, Hungary*

*[4]Institute of Psychology, ELTE, Eötvös Loránd University, Budapest, Hungary*

*[5]UR2NF, Neuropsychology and Functional Neuroimaging Research Unit at CRCN – Center for Research in Cognition and Neurosciences and UNI – ULB Neurosciences Institute, Université Libre de Bruxelles (ULB), Brussels, Belgium*

*[6]Department of General Psychology, Pázmány Péter Catholic University, Budapest, Hungary*

*[7]MTA-PPKE Adolescent Development Research Group, Budapest, Hungary*

*[8]Donders Institute for Brain, Cognition and Behaviour, Radboud University Medical Center, Nijmegen, The Netherlands*

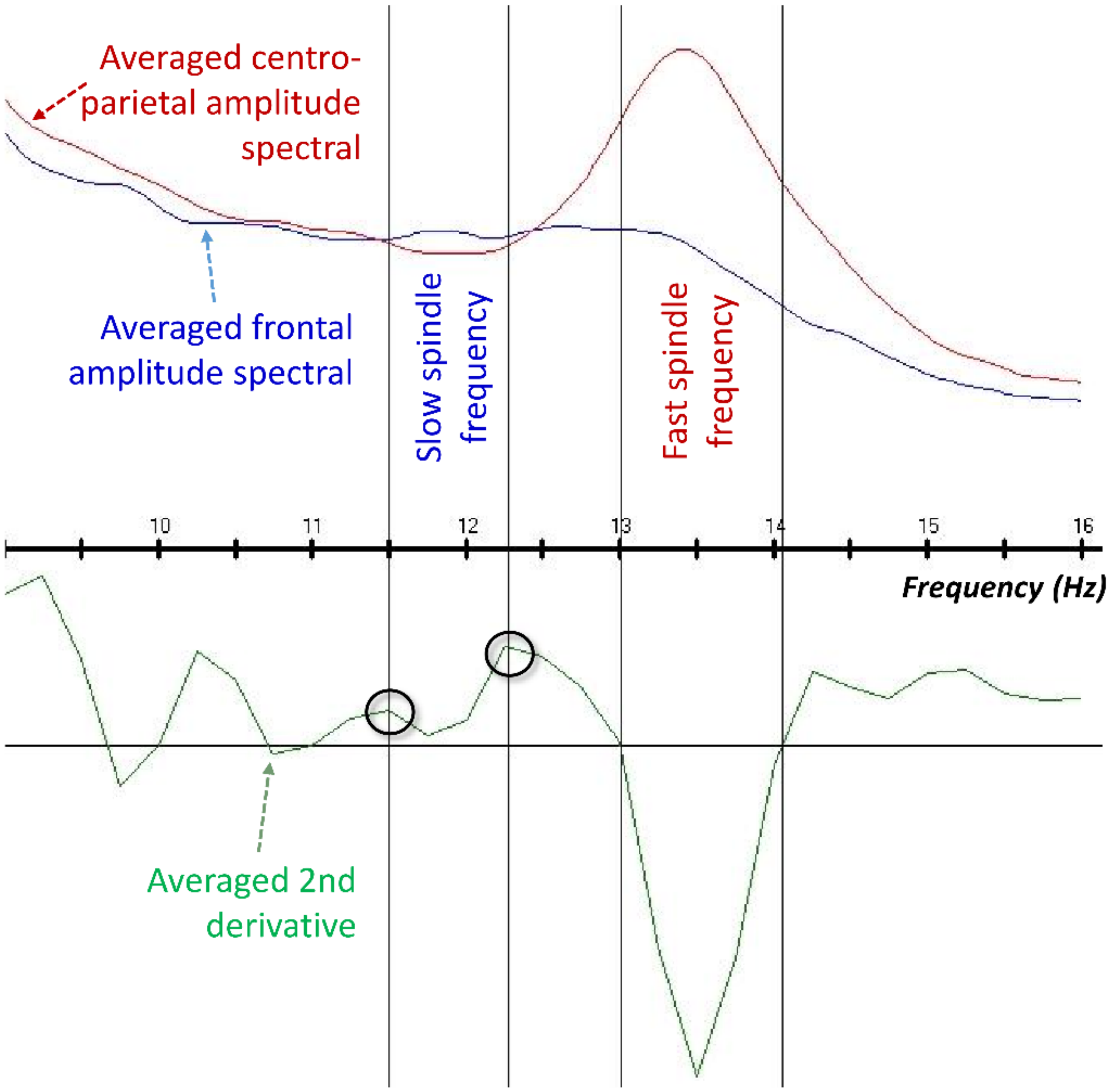


*Supplementary figure 1. Handling the issue of a lack of definitive slow spindle peak in the averaged second derivative of the amplitude spectra. A searching the relative (herewith absolute) maxima of the frontal minus centro-parietal amplitude spectral values was implemented, the slow spindle spectral peaks being defined at the edges of the associated bulge of second-order derivatives (which latter approach but do not reach zero in these cases)*

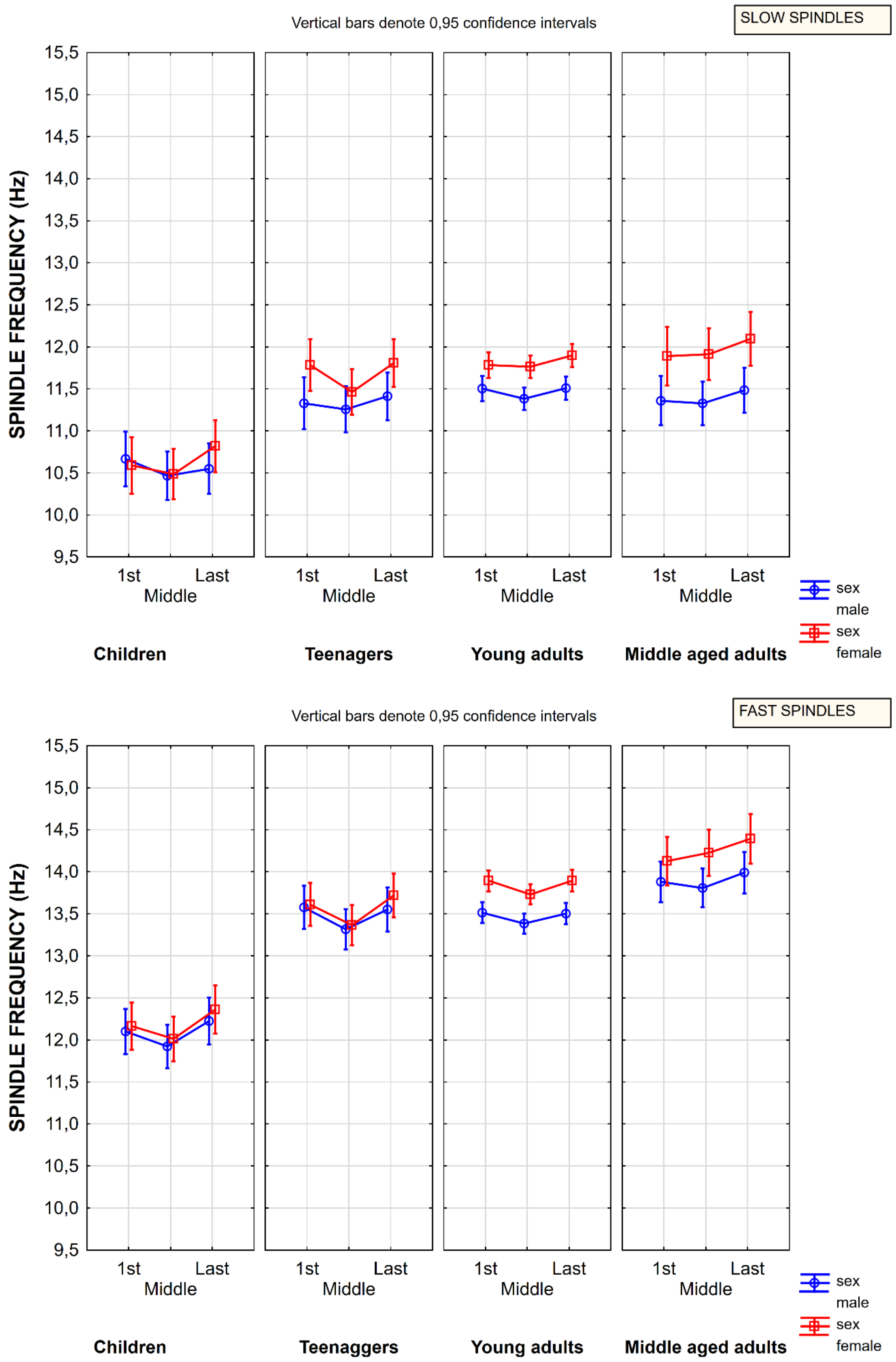


*Supplementary figure 2. Sleep spindle frequency during the first, the middle and the last sleep cycles, with the additional consideration of age and sex.*

Supplementary table 1. Sleep spindle frequencies in the individually determined first, middle and last sleep cycles according to the general linear model*

| Effect | Repeated Measures Analysis of Variance Sigma-restricted parameterization Effective hypothesis decomposition; Std. Error of Estimate: 1.1908 | | | | |
|---|---|---|---|---|---|
| | SS | Degr. of Freedom | MS | F | p |
| Intercept | 150789.4 | 1 | 150789.4 | 106330.8 | 0.000000 |
| sex | 19.7 | 1 | 19.7 | 13.9 | 0.000244 |
| age_group | 287.1 | 3 | 95.7 | 67.5 | 0.000000 |
| sex × age_group | 4.6 | 3 | 1.5 | 1.1 | 0.356934 |
| Error | 344.6 | 243 | 1.4 | | |
| TIME | 7.5 | 2 | 3.7 | 37.4 | 0.000000 |
| TIME × sex | 0.5 | 2 | 0.3 | 2.6 | 0.074094 |
| TIME × age_group | 1.7 | 6 | 0.3 | 2.8 | 0.011684 |
| TIME × sex × age_group | 0.4 | 6 | 0.1 | 0.7 | 0.644557 |
| Error | 48.4 | 486 | 0.1 | | |
| Spindle type | 977.8 | 1 | 977.8 | 2812.2 | 0.000000 |
| Spindle type × sex | 0.7 | 1 | 0.7 | 2.1 | 0.146310 |
| Spindle type × age_group | 17.8 | 3 | 5.9 | 17.0 | 0.000000 |
| Spindle type × sex × age_group | 1.4 | 3 | 0.5 | 1.3 | 0.271605 |
| Error | 84.5 | 243 | 0.3 | | |
| TIME × Spindle type | 0.1 | 2 | 0.0 | 1.1 | 0.326272 |
| TIME × Spindle type × sex | 0.1 | 2 | 0.0 | 0.6 | 0.525841 |
| TIME × Spindle type × age_group | 0.2 | 6 | 0.0 | 1.0 | 0.399486 |
| TIME × Spindle type × sex × age_group | 0.5 | 6 | 0.1 | 2.0 | 0.060602 |
| Error | 19.3 | 486 | 0.0 | | |

**A General Linear Model (GLM) with the between subject factors age group and sex, as well the within subject factors TIME (first, middle and last sleep cycles) and Spindle type (slow and fast) was applied. Middle cycles were determined as follows: the mean of the first and the second cycle in case of 2 complete cycles (2 cases), the second cycle in case of 3 complete cycles, the mean of the second and the third cycles in case of 4 complete cycles and the third cycle in case of 5 complete sleep cycles.*

Supplementary table 2. Additional analysis of nap vs night sleep spindle frequencies in a subsample recorded with the same hardware*

| Effect | Repeated Measures Analysis of Variance Sigma-restricted parameterization Effective hypothesis decomposition; Std. Error of Estimate: 0.6407 | | | | |
|---|---|---|---|---|---|
| | SS | Degr. of Freedom | MS | F | p |
| Intercept | 52060.62 | 1 | 52060.62 | 126819.6 | 0.000000 |
| sex | 14.72 | 1 | 14.72 | 35.8 | 0.000000 |
| Nap_vs_night | 19.13 | 1 | 19.13 | 46.6 | 0.000000 |
| sex × Nap_vs_night | 0.93 | 1 | 0.93 | 2.3 | 0.134746 |
| Error | 74.30 | 181 | 0.41 | | |
| Spindle type | 333.30 | 1 | 333.30 | 2516.9 | 0.000000 |
| Spindle type × sex | 0.01 | 1 | 0.01 | 0.1 | 0.779891 |
| Spindle type × Nap_vs_night | 0.66 | 1 | 0.66 | 5.0 | 0.027291 |
| Spindle type × sex × Nap_vs_night | 0.13 | 1 | 0.13 | 1.0 | 0.319778 |
| Error | 23.97 | 181 | 0.13 | | |

**Nap sleepers (N = 112) and night sleepers (N = 73) did not differ in age (t = 1.26; p = .20) or the recording hardware used for EEG registration (Comlab 32 Digital SleepLab). However, women had faster sleep spindles as compared to males (main effect of sex), whereas nap sleep spindles were of higher frequency than night sleep spindles (main effect of Nap_vs_night). Besides the obvious slow vs fast sleep spindle difference (main effect of Spindle type), slow sleep spindles were characterized by a higher Nap vs night difference than fast sleep spindles (interaction of Spindle type and Nap_vs_night).*
